# Towards feedback control of the cell-cycle across a population of yeast cells

**DOI:** 10.1101/467803

**Authors:** Giansimone Perrino, Davide Fiore, Sara Napolitano, Mario di Bernardo, Diego di Bernardo

## Abstract

Cells are defined by their unique ability to self-replicate through cell division. This periodic process is known as the cell-cycle and it happens with a defined period in each cell. The budding yeast divides asymmetrically with a mother cell generating multiple daughter cells. Within the cell population each cell divides with the same period but asynchronously. Here, we investigate the problem of synchronising the cell-cycle across a population of yeast cells through a microfluidics-based feedback control platform. We propose a theoretical and experimental approach for cell-cycle control by considering a yeast strain that can be forced to start the cell-cycle by changing growth medium. The duration of the cell-cycle is strictly linked to the cell volume growth, hence a hard constraint in the controller design is to prevent excessive volume growth. We experimentally characterised the yeast strain and derived a simplified phase-oscillator model of the cell-cycle. We then designed and implemented three impulsive control strategies to achieve maximal synchronisation across the population and assessed their control performance by numerical simulations. The first two controllers are based on event-triggered strategies, while the third uses a model predictive control (MPC) algorithm to select the sequence of control impulses while satisfying built-in constraints on volume growth. We compared the three strategies by computing two cost functions: one quantifying the level of synchronisation across the cell population and the other volume growth during the process. We demonstrated that the proposed control approaches can effectively achieve an acceptable trade-off between two conflicting control objectives: (i) obtaining maximal synchronisation of the cell cycle across the population while (ii) minimizing volume growth. The results can be used to implement effective strategies to unfold the biological mechanisms controlling cell cycle and volume growth in yeast cells.

## I. Introduction

Attempts to apply engineering principles to biological processes to understand and build new functions in cells have led to a growing interdisciplinary research community blending Synthetic Biology with Control Engineering, aptly named Cybergenetics. Currently synthetic circuits can perform only very basic functions thus having a limited impact in biotechnology and biomedicine. Control engineering is a key discipline that provides the theoretical and methodological tools to engineer complex systems and make sure they behave in the desired way across a range of operating conditions. Application and adaption of established theories and techniques from conventional control engineering have been hampered by the peculiarities of biological systems, such as cell-to-cell variability, metabolic load, cross-talking and practical realizability [1].

Several studies have demonstrated the feasibility of using real-time feedback control approaches to drive simple cellular processes by means of optogenetics or microfluidics approaches [2]–[8], guaranteeing an accuracy that was hitherto unachievable. However, besides proof-of-concepts, the real potential of these approaches has not yet been thoroughly investigated.

Here, we investigate the feasibility of controlling a fundamental and complex mechanism present in each cell: the cell-cycle. This process can be seen as a biological oscillator characterised by a fixed period in each cell, but with different phases across cells. During the cell-cycle, the cell progresses in a sequential and unidirectional manner through different phases (growth, DNA synthesis, mitosis) to make a copy of itself. The control objective we want to achieve is to synchronise the cell cycle across the cell population, thus having all yeast cells budding at the same time. All the available methods in yeast biology do not really “synchronise” the cell population, at least according to the terminology of dynamical system theory, but rather just force each cell in the population to start from the same initial condition. Previous studies addressed the synchronisation problem by considering only an open-loop control strategy where an external periodic input was used to entrain the population of yeast cells, with limited success [9]. Here, we consider the implementation of a feedback control strategy based on an *external* control approach where a computer interfaced with living cells steers their dynamics by means of microfluidics whereas a microscope measures the cells’output.

We first derived a simplified phase-oscillator model of cell-cycle motivated by previous work [9], whose parameters were fitted from experimental data. We used this model both for numerical simulations and for designing control strategies. We designed, numerically simulated and compared three different control strategies, two of which are based on input-output control and one on a model predictive control (MPC) strategy.

We then compared the three strategies against three performance indices that take into account the level of synchronisation across the cell population quantified through the Kuramoto’s order parameter and the cell volume growth during the process.

Our results show the feasibility of applying feedback control strategy to steer a complex biological process such as the cell-cycle and can be instrumental for their implementation in-vivo.

## II. Description of the biological system and characterisation

To address the cell-cycle synchronisation problem, we chose a budding yeast strain whose cell-cycle (shown in Figure 1) can be halted at the G1 phase in the presence of methionine, or allowed to progress to the s phase in the absence of methionine [10].

**Fig. 1.**
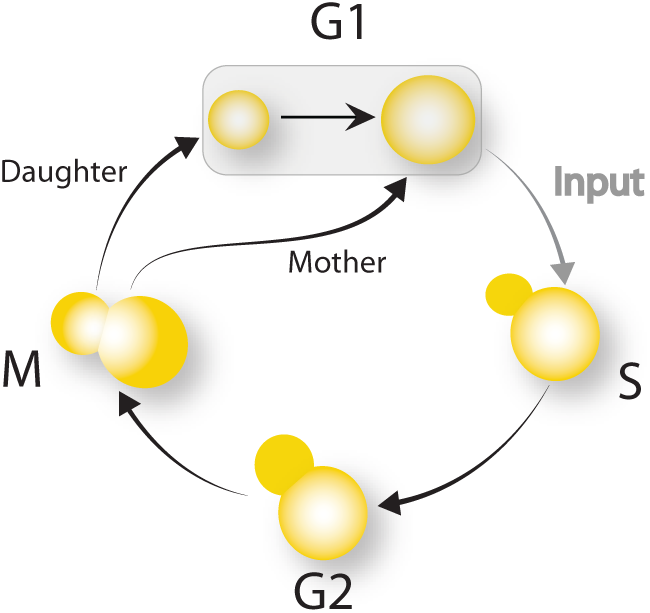
Cell cycle process in a budding yeast manipulated genetically to halt the cell cycle in the G1 phase, unless an external stimulus is provided [10].

### A. Biological system

In the chosen yeast strain, all the endogenous genes encoding the G1 cyclins (*CLN1, CLN2* and *CLN3*) are deleted, thus making the cells unable to progress towards the S phase. To enable control of the cycle progression by changing growth medium, the endogenous *CLN2* gene was inserted downstream of the methionine repressible promoter P_*MET3*_. In this way, the cell-cycle can progress to the S phase only in the absence of methionine. To track the cell-cycle progression, a yellow fluorescent protein (YFP) was inserted downstream of the endogenous *CLN2* promoter. Since the *CLN2* promoter is activated in G1 and repressed in G2, the yellow fluorescence reporter can be used as a proxy of the cell-cycle phase.

### B. Experimental characterisation

We previously implemented a microfluidics experimental platform to enable real-time observation and control of gene expression in yeast cells [8], [11]. We employed this platform to perform a preliminary time-lapse experiment in which cells were grown in the absence of methionine for 18 hours, thus enabling them to cycle. We quantified at single-cell level the YFP fluorescence signal expressed by the endogenous *CLN2* promoter, as well as the area occupied by each cell in the field, as shown in Figure 2A. To reduce the effect of measurement noise, we applied a bandpass Butterworth filter to the fluorescence signals. The filter cut-off frequencies (*f_H_* = 5 · 10^−4^ Hz and *f_L_* = 4 · 10^−5^ Hz) are chosen in the range [30,400] min, as the cell-cycle duration is biologically constrained within this interval.

**Fig. 2.**
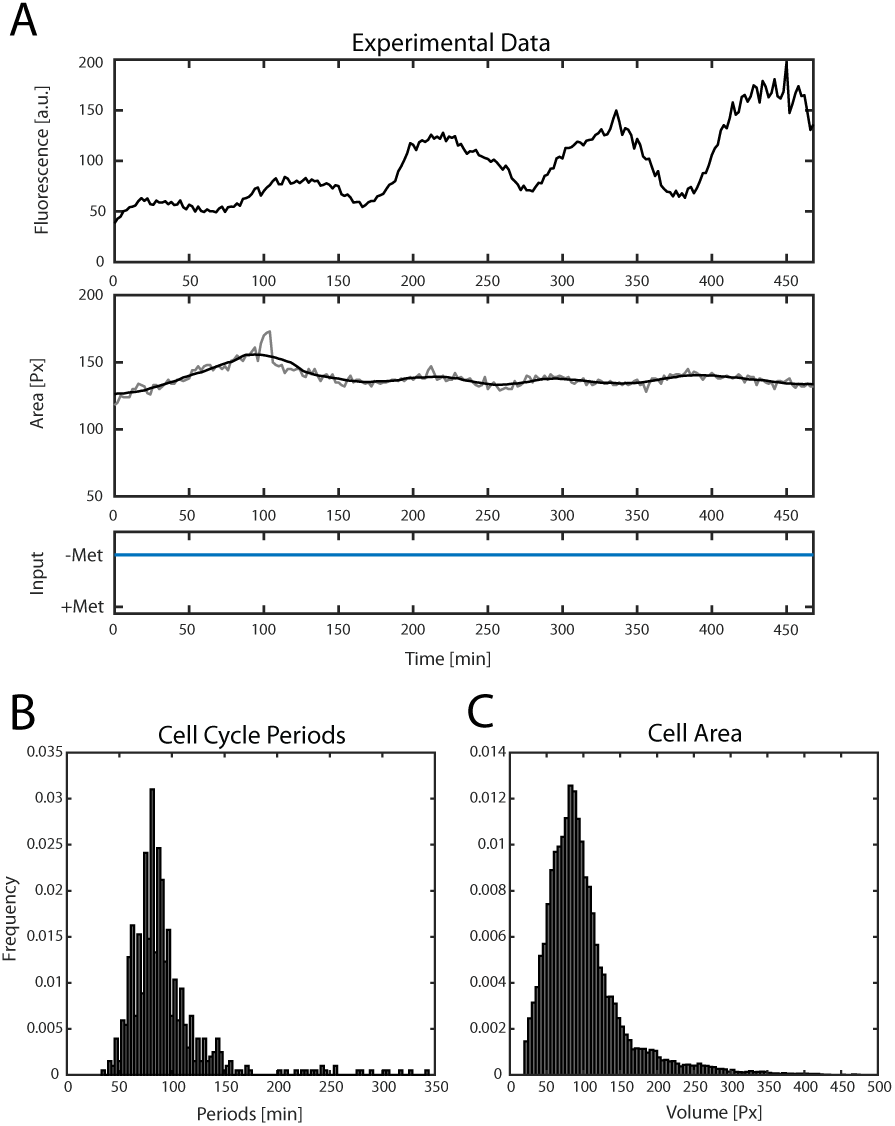
Experimental characterisation of the biological system. (A) Experimental data of a single cell tracked for 500 min. The top panel shows the fluorescence signal measured by a fluorescence microscope at 2 min intervals. The middle panel shows the cell area in pixels. Cells were grown in the absence of methionine, as shown in the bottom panel. (B) Histogram of cell cycle duration across cells. (C) Histogram of cell area across cells.

The measured cell-cycle duration across cells (Figure 2B) can be approximated by a normal distribution 𝓝 (*μ*, *σ*^2^) with mean *μ* = 84.47 min and standard deviation *σ* = 19.33 min. As expected, the mean value *μ* is in line with previous observations [10]. We noted a high value for the standard deviation *σ*, which may be explained by the presence of cells in the field that are not cycling despite the activation of the endogenous gene *CLN2*. The estimated cell areas, which are proportional to the volume, are distributed as a log-normal distribution with mean *μ* = 4.48 px and standard deviation *σ* = 0.4 px (Figure 2C).

### C. Model derivation

To model the cell-cycle, we exploited the concept of phase reduction [12], [13]. We assumed that the cell-cycle can be described as a dynamical system of the general form:

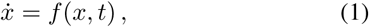

where *x* ∊ ℝ^*n*^ is the state vector and *t* ∊ ℝ is time. If (1) has an exponentially stable limit cycle *γ* ⊂ ℝ^*n*^ with period *T_d_*, then (1) is an oscillator that, according the phase reduction method, can be modelled as a dynamical phase oscillator

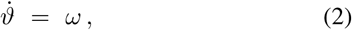

where *ϑ* is the phase of the oscillator on the unit circle 𝕊^1^ and
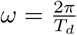
is the angular frequency.

Let *ϑ_c_* be the cell-cycle phase at budding, i.e. the phase at which the G1/S transition occurs. As the cell cycle is always in G1 in the presence of methionine, the phase dynamics can be described as:

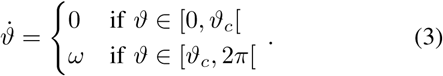

Thus, the phase linearly increases with rate *ω* only when the cell is in either S, G2 or M. As the cell-cycle is coupled to cell growth [9], we consider also cell volume dynamics. To this aim, we assume that a cell grows exponentially only during the G1 phase. Thus, the volume growth can be described as:

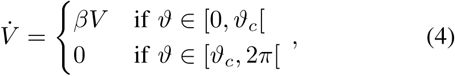

where *β* > 0 is the volume growth rate. We assume that cell division occurs at *ϑ* = 2*π*. At each division event, the phase *ϑ* is reset to *ϑ* = 0.

Let *u* ∈ {0, 1} be the external trigger input to the system. We assume that, if *u* = 1, the cells are growing in the absence of methionine; conversely, if *u* = 0, they are growing in the presence of methionine. We then model the phase transition from G1 to S as the following reset condition:

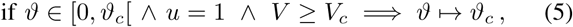

where *V_c_* is the critical volume, defined as the minimum amount of volume needed to have the cell cycle progress towards the S phase.

## III. Problem statement

Let *N* ∈ ℕ be the number of cells in the population. Consider each cell as an individual agent whose cell-cycle progression is mathematically described by the model composed by (3) and (4), together with the reset condition (5).

The control aim is to synchronise the cell-cycle across the cell population. As the duration of the cell-cycle is coupled to the volume growth, a desirable feature of the controller is to avoid excessive volume growth, which can be lethal for the cell.

Synchronisation in a population of oscillators can be quantified by means of the Kuramoto order parameter *R* [13] defined as the magnitude of the complex number

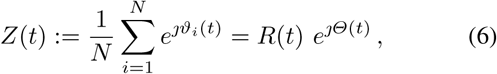

where *ϑ_i_* is the phase of cell *i*. *R* ∈ [0, 1] represents the mean phase coherence, an index to evaluate the synchronisation among a population of oscillators. When *R* is equal to 1, all cells are synchronized onto the same phase.

Cell volume is quantified by measuring the average volume in the population:

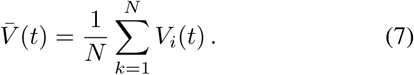

To evaluate the performance of the control algorithms, we introduce two cost functions:

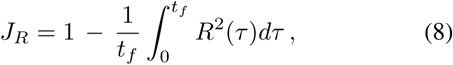

so that all cells are synchronized over the whole time interval when *J_R_* is equal to zero; and:

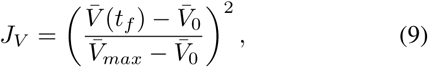

with *J_V_* [0, 1],
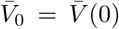
and
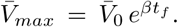
Here
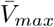 is the maximum value of the average volume that would be reached if all cells exponentially grew for all *t* ∈ [0, *t_f_*]. Notice that volume growth is minimized over the time interval being considered if *J_V_* is rendered as close as possible to 0.

The control problem can thus be stated as follows:

### Problem 1

Given a population of 𝓝 individual cells, which are described by (3), (4), (5), compute the control input *u* that minimises *J_R_* and, possibly, *J_V_*.

### A. Numerical simulations

All in-silico simulations were performed using the Matlab `ode45` solver with event detection routines to accurately detect cell division at *ϑ_i_* = 2*π* and the hitting of the critical volume *V_i_* = *V_c_* for cells with initial volume *V_i_* < *V_c_*.

We considered a synthetic population consisting of 𝓝 = 100 cells with random initial conditions. Specifically, the initial volumes *V_i_*(0) were drawn from the interval [0.8 *V_c_*, 1.2 *V_c_*] and the initial phases *ϑ_i_*(0) from the interval [0, 2*π*], both with uniform distributions, and setting *ϑ_i_* to 0 if *V_i_* < *V_c_*. Moreover we set
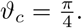

The volume growth rate is chosen as *β* = 0.0083 [a.u.]min^−1^ and the critical volume as *V_c_* = 1000 [a.u.], following the same choice reported in [9].

Due to the high standard deviation in the cell cycle periods *T_i_* identified from the experimental data, we considered also an additional case with a lower standard deviation (hence a lower value of the coefficient of variation *CV* = *σ/μ*, where *μ* is the nominal cell cycle period *T*). Specifically:

- High variability case (*CV* = 22.88%): *T_i_* drawn from the normal distribution with parameters *μ* = 84.47 min and *σ* = 19.33 min, winsorizing samples outside [40, 138] min.
- Low variability case (*CV* = 10%): *T_i_* drawn from the normal distribution with parameters *μ* = 84.47 min and *σ* = 8.447 min, winsorizing samples outside [67, 101] min.

All simulations were stopped at *t_f_* = 6*T*.

## IV. Control algorithms

We designed three control algorithms to solve Problem 1, that is to achieve the synchronisation of the yeast cell cycle, and concurrently the minimisation of the volume growth in heterogeneous populations of cells. The first control algorithm we present, named *Stop&Go control*, is trivial and hence it is used as benchmark to assess the performance of the other two more advanced control algorithms. The second control algorithm, that we named *Threshold control*, is an empirical control strategy. The third control algorithm is a *model predictive control* (MPC), which is widely exploited for external in-silico control of biological systems [6]–[8], [14].

### A. Stop&Go control

The Stop&Go control algorithm consists into applying a control trigger *u* = 1 every time *all* cells are at *ϑ_i_* = 0, that is they have stopped at the G1 phase:

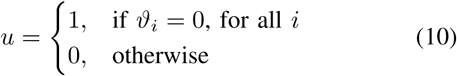

At steady-state, that is after the first trigger, the control signal *u*(*t*) becomes periodic with period
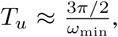
where *ω*_min_ is the angular frequency of the slowest cell in the population. This algorithm does not attempt to minimise the cell volume growth. Figure 3(a) depicts the phase *ϑ_i_*(*t*) and volume *V_i_*(*t*) normalised by the critical volume *V_c_* of the cells in the population together with the control input generated by this algorithm. The time evolution of the mean phase coherence *R*(*t*) and the normalised average population volume
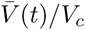
are reported in Figure 3(b) where they are averaged over 10 trials with random initial conditions.

**Fig. 3.**
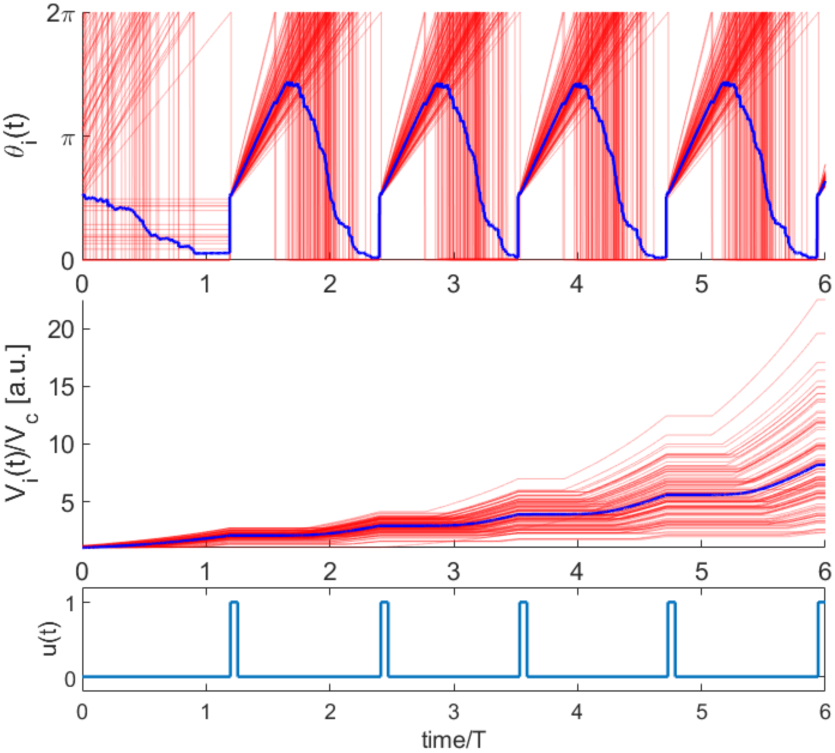
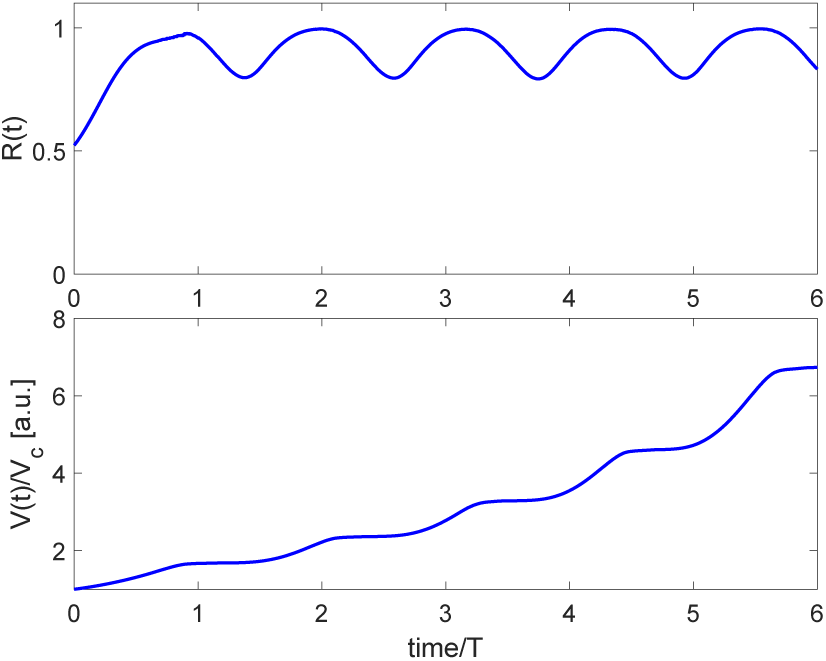
Stop&Go control: High-variability of *T_i_* (*CV* = 22.88%). (a) Time evolution of *ϑ_i_*(*t*) and 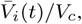 for one simulation trial, are reported in red colour. Their average values over the whole population are reported in blue. The corresponding control signal *u*(*t*) generated by the Stop&Go algorithm is reported in the bottom panel. (b) Time evolution of *R*(*t*) and 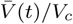 averaged over 10 simulation trials with random initial conditions.

### B. Threshold control

The Threshold control algorithm is a heuristic control strategy where a trigger *u* = 1 is applied only when there are no cells with phase *ϑ_i_* in a certain interval 𝓘(*k*), that is

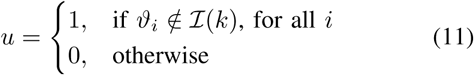

This interval is defined as

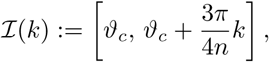

where *n* is the *first control parameter* and represents the number of sub-intervals in which the sector
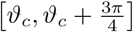
is divided, as shown in blue in Figure 4. The right endpoint of the interval 𝓘(*k*) defines the *current threshold ϑ_u_*(*k*) =
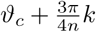
whose values grows with *k* ∈ {1, …, *n*, 2*n*}. The evolution in time of *k* is defined as

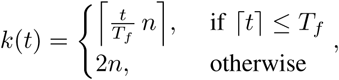

where *T_f_* < *t_f_* is a *second control parameter* that represents the first time instant at which we want *k* = 2*n*. Therefore, at *t* = *T_f_* we have *ϑ_u_*(2*n*) = 2*π*, as
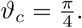
Notice that when this condition is met, the control laws (10) and (11) become equivalent and therefore this control algorithm behaves as the Stop&Go control.

**Fig. 4.**
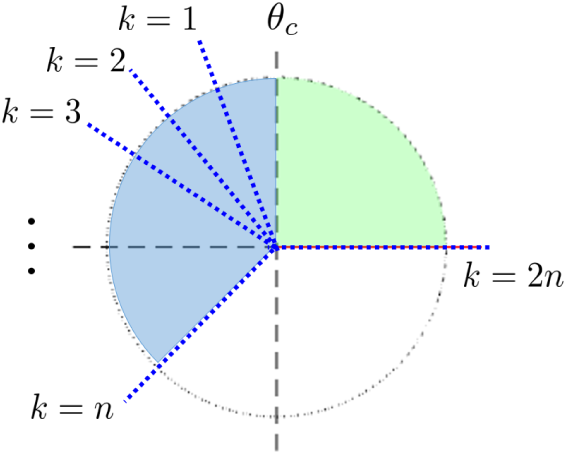
Illustration of the evolution of the threshold *ϑ_u_* in Threshold control algorithm. The phase interval
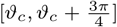
(depicted in blue) is divided into *n* equal subintervals. As *k*(*t*) grows in time from 1 to 2*n*, the interval 𝓘(*k*) with *ϑ_u_*. The G1 phase is depicted in green, corresponding to the phase interval [0, *ϑ_c_*]. Cell phases are assumed to positively evolve in the anticlockwise direction.

Were the algorithm to be applied to a single cell, then the number of control triggers, say *N_u_* over a certain time interval [0, *t_f_*] could be analytically computed as:

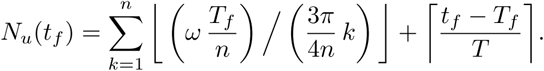

However, due to the heterogeneity of cell-cycle duration across cells this number represents only an upper bound on the actual number of control triggers generated by the algorithm.

To limit volume growth (*J_V_*), this algorithm is complemented with a time-dependent control. Specifically, even when the condition (11) on the phases is not satisfied, a control trigger is applied if the last trigger was applied more than
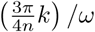
minutes ago, where *ω* is the nominal cell cycle angular frequency, that is *ω* = 2*π/T*. This allows to avoid unnecessary volume growth due to wasted time waiting for “slow” cells to reach the current threshold *ϑ_u_*(*k*).

Figure 5(a) shows the response of the cell population to the control input generated by the Threshold algorithm with control parameters (*n*, *T_f_*) = (8, 2*T*). The evolution of *R*(*t*) and 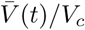 averaged over 10 simulation trials with random initial conditions and selected values of the control parameters (*n*, *T_f_*) are reported in Figure 5(b).

**Fig. 5.**
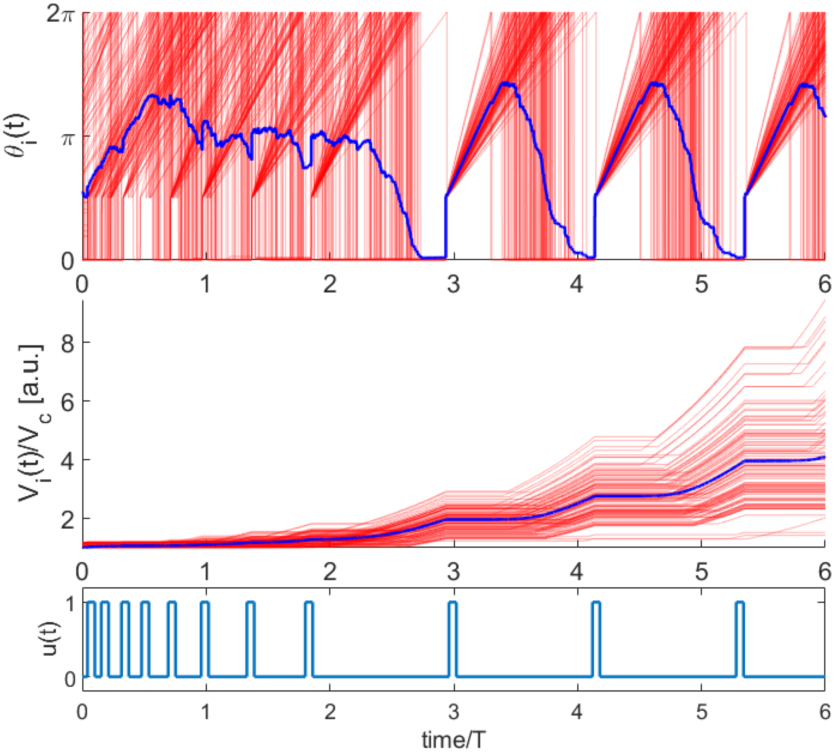
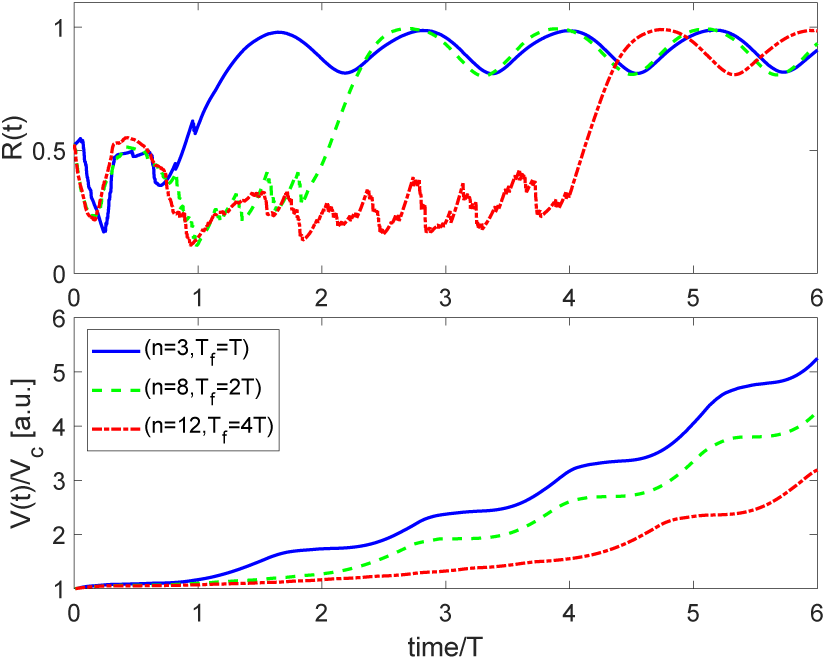
Threshold control: High-variability of *T_i_* (*CV* = 22.88%). (a) Time evolution of *ϑ_i_*(*t*) and 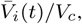 for one simulation trial, are reported in red. Average values over the whole population are reported in blue. The corresponding control signal *u*(*t*) generated by the Threshold control algorithm with parameters *n* = 8, *T_f_* = 2*T* is reported in the bottom panel. (b) Time evolution of *R*(*t*) and 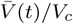 averaged over 10 simulation trials with random initial conditions for three different sets of control parameters: (*n*, *T_f_*) = (3, *T*) in solid blue lines, (*n*, *T_f_*) = (8, 2*T*) in green dashed lines, (*n*, *T_f_*) = (12, 4*T*) in red dotted lines.

### C. Model Predictive Control

The MPC algorithm consists in solving an open-loop optimal control problem repeatedly over a receding horizon [15]. This means that at each iteration of the algorithm, the solution of the optimal open-loop control problem gives an optimal input that minimises a cost function over a finite prediction horizon *T_p_*. The optimal input is applied over a finite control horizon *T_c_* ≤ *T_p_*, hence discarding the remaining part of the computed optimal control action. Then, the optimisation is repeated again.

To address the problem of minimising the two conflicting performance indices *J_R_* and *J_V_*, we chose the cost function

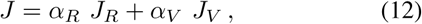

which combines both *J_R_* and *J_V_*, defined in (8) and (9), with *t_f_* = *T_p_*, by assigning a different weight to each index function (*weighting method*) [16]. The choice of the weights *α_R_* and *α_V_*, with *α_R_* ≥ 0, *α_V_* ≥ 0, *α_R_* + *α_V_* = 1, affects the overall performance of the algorithm, hence they are considered as further control parameters, and represent a trade-off between the two different control objectives.

To reduce the computational complexity of the optimal control problem, the control input *u* ∈ 𝓤 is assumed to be a finite sequence of triggers. Defining *P* ∈ ℕ as the maximum number of triggers that may be applied in a finite prediction horizon, then the time interval occurring between two consecutive triggers is
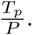
The feasible set 𝓤 is defined as the set of all admissible control sequences. Considering only the finite prediction horizon *T_p_*, the set 𝓤 is composed by 2^*P*^ possible combinations of triggers. Thus, the optimal control problem is solved by minimising the performance index *J* over the prediction horizon *T_p_*. The optimisation is achieved by exploring all the possible combinations of sequence of triggers.

For the numerical analysis, we set the length of both the prediction horizon *T_p_* and the control horizon *T_c_* equal to the nominal cell-cycle duration *T*, and the number of triggers *P* equals to 6. To assess the feasibility of the control algorithm, a set of simulations was carried out by varying the weights *α_R_* and *α_R_*. Figure 6(a) shows the response of the cell population to the control input generated by the MPC algorithm with control parameters (*α_R_*, *α_V_*) = (0.4, 0.6). Examples of three time evolution of the mean phase coherence *R*(*t*) and the normalised average population volume 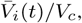 both averaged over 10 trials with random initial conditions are shown in Figure 6(b).

**Fig. 6.**
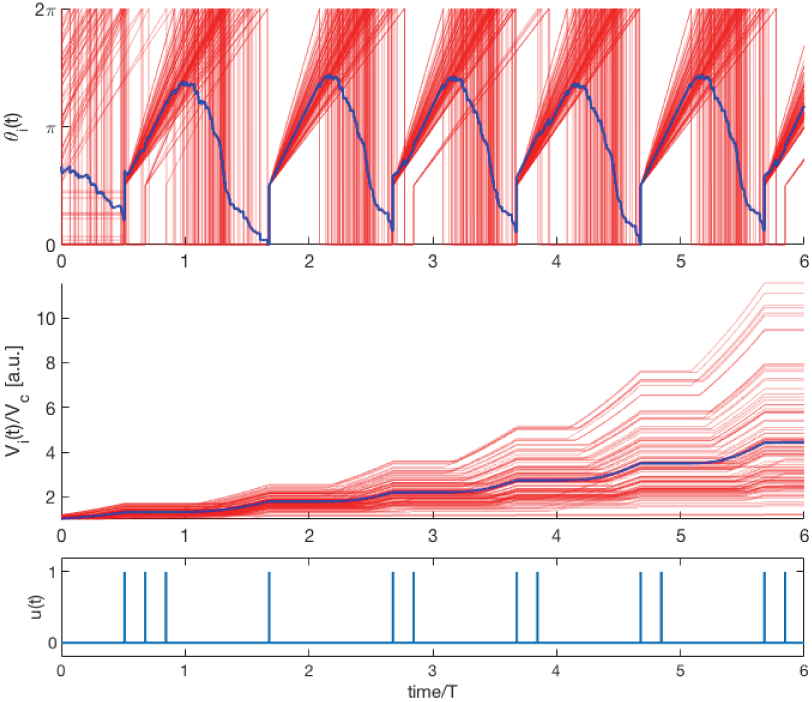
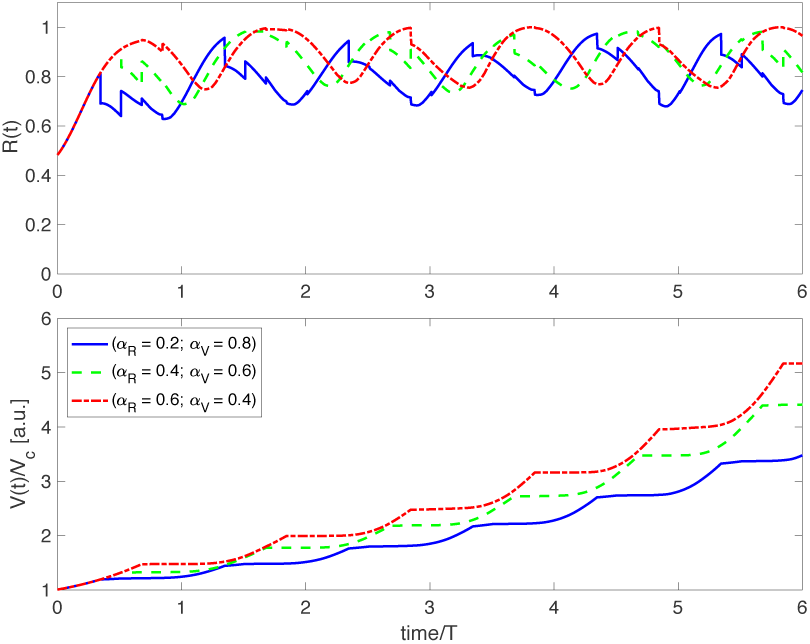
MPC: High-variability of *T_i_* (*CV* = 22.88%). (a) Time evolution of *ϑ_i_*(*t*) and 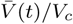 for one simulation trial, are reported in red colour. Their average values over the whole population are reported in blue. The corresponding control signal *u*(*t*) generated by the MPC algorithm with weight factors (*α_R_*, *α_V_*) = (0.4, 0.6) is reported in the bottom panel. (b) Time evolution of *R*(*t*) and 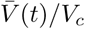 averaged over 10 simulation trial with random initial conditions for three different sets of weight factors (*α_R_*, *α_V_*) = (0.2, 0.8) in solid blue lines, (*α_R_*, *α_V_*) = (0.4, 0.6) in gree dashed lines, (*αR, αV*) = (0.6, 0.4) in red dotted lines.

### D. Comparative analysis

We performed a comparative analysis the three control algorithms across several simulated scenarios. For each scenario, we computed the indices *J_R_*, *J_V_*, and the total number of triggers applied during the simulation. Results obtained averaging over 10 trials with random initial conditions are presented in Table I.

**TABLE I.**
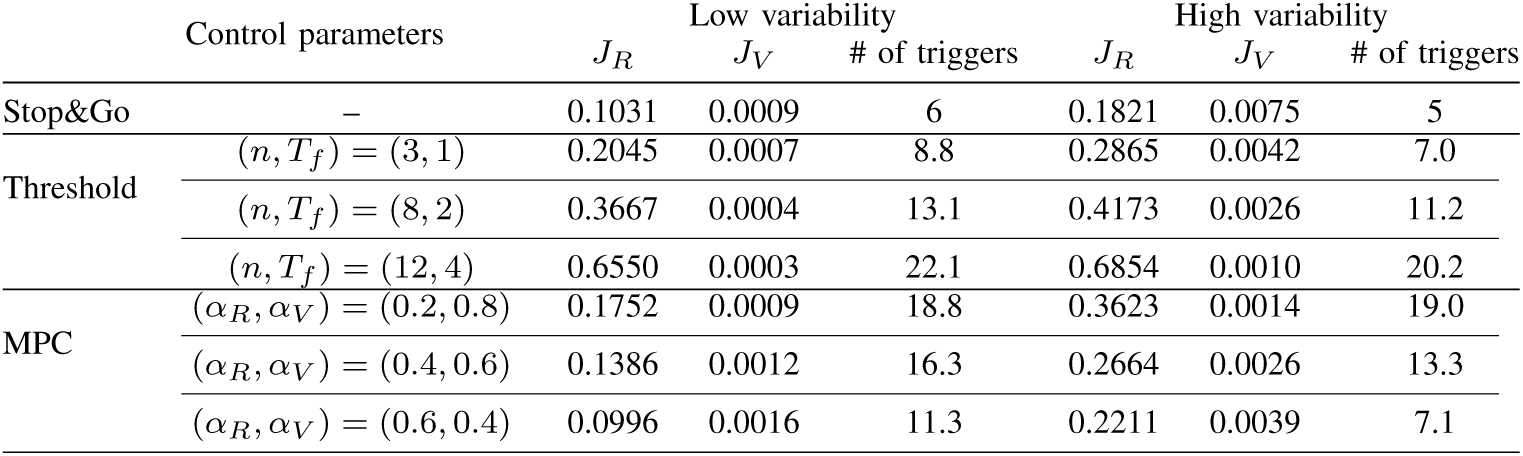
NUMERICAL COMPARISON OF THE CONTROL ALGORITHMS.

#### a) Stop&Go control

Although this control algorithm is very simple and can guarantee a high level of synchronisation (*J_R_* equal to 0.1821 and 0.1031 for high and low cell-to-cell variability, respectively) reaching a regime after about one nominal period *T* of the cell cycle, it has the drawback of causing excessive volume growth (*J_V_* equal to 0.0075 and 0.0009 for high and low cell-to-cell variability, respectively). Indeed, faster cells, i.e. those having a smaller cycle period *T_i_*, reach phase 2*π* before the other slower cells and therefore they are stuck there growing until the next control trigger is applied, thus yielding an unnecessary volume increase.

#### b) Threshold control

This control algorithm guarantees a good level of synchronisation and it contains excessive volume growth with respect to the Stop&Go control. However, a trade-off between *J_R_* and *J_V_* exists in the tuning of the control parameters (*n*, *T_f_*). Specifically, low values of *T_f_* give lower values of *J_R_* but higher values of *J_V_*, and viceversa for high values of *T_f_*. On the other hand, increasing *n* gives lower values of *J_V_* but it does not result in significant reduction of *J_R_*.

#### c) MPC

This control algorithm reaches a high level of synchronisation, limiting at the same time the volume growth. Among the control algorithms, the MPC achieves the best trade-off between the indices *J_R_* and *J_V_*. One drawback of the algorithm could be the computational effort needed to solve the optimal control problem in real-time. However, the assumption on the finite sequence of triggers reduces the computational time, allowing the solution of the optimal control problem to be found in few seconds.

## V. Conclusions

We have presented a theoretical and experimental framework to address the cell-cycle synchronisation problem in budding yeast in the context of a real biological scenario. We characterised experimentally a yeast strain whose cell-cycle can be controlled experimentally by changing growth medium. We then derived a phase-oscillator model to describe the cell-cycle phase, as well as the volume growth dynamics. We demonstrated in-silico the feasibility of synchronizing the cell cycle across a cell population, while containing at the same time the volume growth. To this end, we devised and implemented three control algorithms: *Stop&Go control*, *Threshold control* and *MPC*. A numerical analysis was performed to assess and compare the three strategies.

Taken together all the numerical results provide evidence for the success of the proposed control approaches in living yeast cells. Nevertheless, several issues remain open both in terms of control performance and experimental implementation. Indeed, model uncertainty and parametric variability could affect the control performance, thus adaptive and robust control strategies are required. To this aim, we aim to improve the control strategies by choosing adaptively the control algorithms’ parameters, e.g. *α_R_* and *α_V_* of MPC algorithm, or resorting to alternative heuristic strategies such as reinforcement learning control.

Finally, ongoing work is aimed at implementing and validating in-vivo the proposed control strategies which will be presented elsewhere.

## Acknowledgement

The authors would like to thank Prof. Sahand Jamal Rahi (EPFL, Switzerland) for providing the budding yeast strain used to conduct this study. The authors also wish to acknowledge support from the research project COSY-BIO (Control Engineering of Biological Systems for Reliable Synthetic Biology Applications) funded by the European Union’s Horizon 2020 research and innovation programme under grant agreement No 766840.

